# *Streptococcus pneumoniae*, *S. pyogenes*, and *S. agalactiae* membrane phospholipid remodeling in response to human serum

**DOI:** 10.1101/2021.01.28.428653

**Authors:** Luke. R. Joyce, Ziqiang Guan, Kelli L. Palmer

## Abstract

*Streptococcus pneumoniae*, *S. pyogenes* (Group A *Streptococcus*; GAS), and *S. agalactiae* (Group B *Streptococcus*; GBS) are major etiological agents of diseases in humans. The cellular membrane, a crucial site in host-pathogen interactions, is poorly characterized in streptococci. Moreover, little is known about whether or how environmental conditions influence their lipid compositions. Using normal phase liquid chromatography coupled with electrospray ionization mass spectrometry, we characterized the phospholipids and glycolipids of *S. pneumoniae*, GAS, and GBS in routine undefined laboratory medium, streptococcal defined medium, and, in order to mimic the host environment, defined medium supplemented with human serum. In human serum-supplemented medium, all three streptococcal species synthesize phosphatidylcholine (PC), a zwitterionic phospholipid commonly found in eukaryotes but relatively rare in bacteria. We previously reported that *S. pneumoniae* utilizes the glycerophosphocholine (GPC) biosynthetic pathway to synthesize PC. Through substrate tracing experiments, we confirm that GAS and GBS scavenge lysoPC, a major metabolite in human serum, thereby using an abbreviated GPC pathway for PC biosynthesis. Furthermore, we found that plasmanyl-PC is uniquely present in the GBS membrane during growth with human serum, suggesting GBS possesses unusual membrane biochemical or biophysical properties. In summary, we report cellular lipid remodeling by the major pathogenic streptococci in response to metabolites present in human serum.

## Introduction

Streptococci are Gram-positive bacteria that natively colonize humans in niches including the oral cavity, the nasopharynx, and the genitourinary tract (1). Three species of streptococci, *Streptococcus pneumoniae*, *S. pyogenes* (Group A *Streptococcus*; GAS), and *S. agalactiae* (Group B *Streptococcus*; GBS), are considered major human pathogens, resulting in over 1.5 million deaths each year around the world (2,3). These pathogens cause a wide range of diseases of varying severity in all age groups, including streptococcal pharyngitis, soft tissue infections, pneumonia, meningitis, bacteremia, and necrotizing fasciitis (3–10).

The surface of the streptococcal cell is a critical interface for host interactions. Many studies into the pathogenicity of streptococci have focused on virulence factors including secreted proteins, surface polymers and adhesins, and mechanisms for sensing the extracellular environment (11–13). In contrast, little is known about the cellular membrane composition and dynamics of streptococci. Earlier studies mainly utilized thin layer chromatography (TLC) or column separation coupled with paper chromatography to study membrane lipids extracted from cultures in routine laboratory medium (14–19). However, these methods lack molecular specificity and sensitivity for the comprehensive characterization of membrane lipids (20). The phospholipids phosphatidylglycerol (PG) and cardiolipin (CL), and the glycolipids monohexosyldiacylglycerol (MHDAG) and dihexosyldiacylglycerol (DHDAG) were previously detected in all three streptococcal species (14–19,21–23).

We recently utilized normal phase liquid chromatography coupled with electrospray ionization mass spectrometry (NPLC-ESI/MS) to characterize the lipid membranes of the Mitis group streptococcal species *S. pneumoniae*, *S. mitis*, and *S. oralis*, and described for the first time the presence of the zwitterionic phospholipid phosphatidylcholine (PC) in these organisms (24,25). We further showed that the biosynthesis of PC in these organisms occurs through the rare glycerophosphocholine (GPC) pathway which relies upon scavenging of GPC and/or lysophosphatidylcholine (lysoPC) from the growth medium (24).

Here, we utilized NPLC-ESI/MS to characterize the membrane lipids of the three major streptococcal pathogens. To mimic the human host environment and to assess the effects of human serum on the membrane lipid compositions, the streptococci were cultured in defined medium with or without human serum supplementation. We show that: 1) human serum provides the substrates required for *S. pneumoniae* to synthesize PC; 2) the GAS and GBS scavenge the human metabolite lysoPC to synthesize PC via an abbreviated GPC pathway; and, 3) plasmanyl-PC (pPC) is uniquely present in the GBS membrane during culture with human serum.

## Material and Methods

### Bacterial strains, media, and growth conditions

Culture media and bacterial strains used in this study are shown in Table 1 and Table S1, respectively. Routine laboratory culture conditions for each species were 37°C for GBS, and 37°C and 5% CO_2_ for *S. pneumoniae* and GAS. Rich culture media used were Todd-Hewitt Broth for GBS and Todd-Hewitt Broth supplemented with yeast extract at 0.5% w/v for *S. pneumoniae* and 0.2% w/v for GAS. Streptococcal chemically defined medium (26) was diluted from stock as described (27) with 1% glucose, for GAS and GBS. For *S. pneumoniae*, defined medium was supplemented with 0.5 mM choline (25). Where appropriate, defined medium was supplemented with 5% v/v human serum (Sigma-Aldrich), 100 μM GPC (Sigma-Aldrich), or 100 μM lysoPC 20:0 (Avanti Polar Lipids). See Supplemental Text S1 for growth curve methodology.

**Table 1.** Major lipids detected in *S. pneumoniae*, GAS, and GBS.

| Species | Strain | Medium<sup>1</sup> | Detection of<sup>2</sup> |  |  |  |  |  |  |  |  |  |  |
| --- | --- | --- | --- | --- | --- | --- | --- | --- | --- | --- | --- | --- | --- |
|  |  |  | DAG | MHDAG | DHDAG | PA | PG | CL | C<sub>55</sub>-P | Lys-PG | L<br>G<br>D | PC | pPC |
| *S. pneumoniae*<sup>3</sup> | D39 | THB+0.5Y | + | + | + | + | + | + | + | - | - | + | - |
|  |  | DM | + | + | + | + | + | + | + | - | - | - | - |
|  |  | DM-HS | + | + | + | + | + | + | + | - | - | + | - |
|  | TIGR4 | THB+0.5Y | + | + | + | + | + | + | + | - | - | + | - |
|  |  | DM | + | + | + | + | + | + | + | - | - | - | - |
|  |  | DM-HS | + | + | + | + | + | + | + | - | - | + | - |
| *S. pyogenes*<br>(GAS) | NZ131 | THB+0.2Y | + | + | + | + | + | + | + | - | - | - | - |
|  |  | DM | + | + | + | + | + | + | + | - | - | - | - |
|  |  | DM-HS | + | + | + | + | + | + | + | - | - | + | - |
|  | MGAS<br>315 | THB+0.2Y | + | + | + | + | + | + | + | - | - | - | - |
|  |  | DM | + | + | + | + | + | + | + | - | - | - | - |
|  |  | DM-HS | + | + | + | + | + | + | + | - | - | + | - |
| *S. agalactiae*<br>(GBS) | COH1 | THB | + | + | + | + | + | + | + | + | + | - | - |
|  |  | DM | + | + | + | + | + | + | + | + | + | - | - |
|  |  | DM-HS | + | + | + | + | + | + | + | + | + | - | + |
|  | A909 | THB | + | + | + | + | + | + | + | + | + | - | - |
|  |  | DM | + | + | + | + | + | + | + | + | + | - | - |
|  |  | DM-HS | + | + | + | + | + | + | + | + | + | - | + |

<sup>1</sup>Todd-Hewitt Broth (THB); THB supplemented with 0.5% yeast extract (THB+0.5Y); THB supplemented with 0.2% yeast extract (THB+0.2Y); Streptococcal defined medium (DM), Streptococcal defined medium supplemented with 5% v/v human serum (DM-HS).
<sup>2</sup>Abbreviations and denotations: + , detected; -, undetected; DAG, diacylglycerol; MHDAG, monohexosyldiacylglycerol; DHDAG, dihexosyldiacylglycerol; PA, phosphatidic acid; PG, phosphatidylglycerol; CL, cardiolipin; C<sub>55</sub>-P, undecaprenyl phosphate; Lys-PG, lysyl-phosphatidylglycerol; LGD, lysyl-glucosyl-DAG; PC, phosphatidylcholine; pPC, plasmalogen-PC.
<sup>3</sup>*S. pneumoniae* lipid profiles in THB+0.5Y and defined medium were originally described in Joyce LR *et al* 2019 (25).

### GAS and GBS culture for lipidomic analysis

Single colonies of GAS and GBS were inoculated into 15 mL rich medium and incubated overnight as described above, until pelleting and storage for lipidomic analysis described below. For defined medium analysis, single colonies were cultured in defined medium overnight, diluted into 15 mL pre-warmed defined medium supplemented with or without 5% v/v human serum to an OD_600nm_ of 0.05, and incubated for 8 h before pelleting. Cultures were performed in biological triplicate.

### *S. pneumoniae* culture for lipidomic analysis

Single colonies of *S. pneumoniae* were inoculated into 6 mL rich medium, serially diluted, and incubated overnight as described. Cultures in early exponential phase were inoculated into 15 mL pre-warmed rich medium and cultured until early stationary phase. Overnight defined medium cultures were inoculated directly from freezer stocks and serially diluted for overnight growth. Early exponential phase cultures were inoculated into defined medium supplemented with or without 5% v/v human serum at an OD_600nm_ of 0.05 and incubated for 8 h until pelleting. Cultures were performed in biological triplicate.

### GPC and lysoPC supplementation experiments

Overnight defined medium cultures of GAS and GBS were diluted to a starting OD_600nm_ 0.05 in 15 mL defined medium supplemented with either 100 μM GPC or 100 μM lysoPC (20:0) and incubated for 8 h as described above. Cultures were pelleted and stored for lipidomic analysis. Cultures were performed in biological triplicate.

### Lipidomics

Acidic Bligh-Dyer lipid extractions were performed as described (25,28). Normal-phase LC-ESI/MS was performed on an Agilent 1200 quaternary LC system equipped with an Ascentis silica high-performance liquid chromatography (HPLC) column (5 μm; 25 cm by 2.1 mm; Sigma-Aldrich) as described previously (25,28,29). See Supplemental Text S1 for more information.

## Results

### Pathogenic streptococci remodel their membrane phospholipid composition in response to human serum

The major phospholipids detected by normal-phase LC-ESI/MS for each streptococcal species and strain used in this study, when cultured in the rich, undefined laboratory medium Todd-Hewitt Broth (supplemented with yeast extract where appropriate), and in streptococcal defined medium supplemented with or without 5% v/v human serum, are shown in Table 1. The biosynthetic pathways for these lipids are shown in Figure 1. Growth curves for streptococci cultured with or without 5% v/v human serum are shown in Supplemental Figure S1A-F. Human serum supplementation did not significantly alter the growth of the bacteria, except during late exponential and early stationary phase of *S. pneumoniae* D39 (Supplemental Figure S1A).

**Figure 1.**
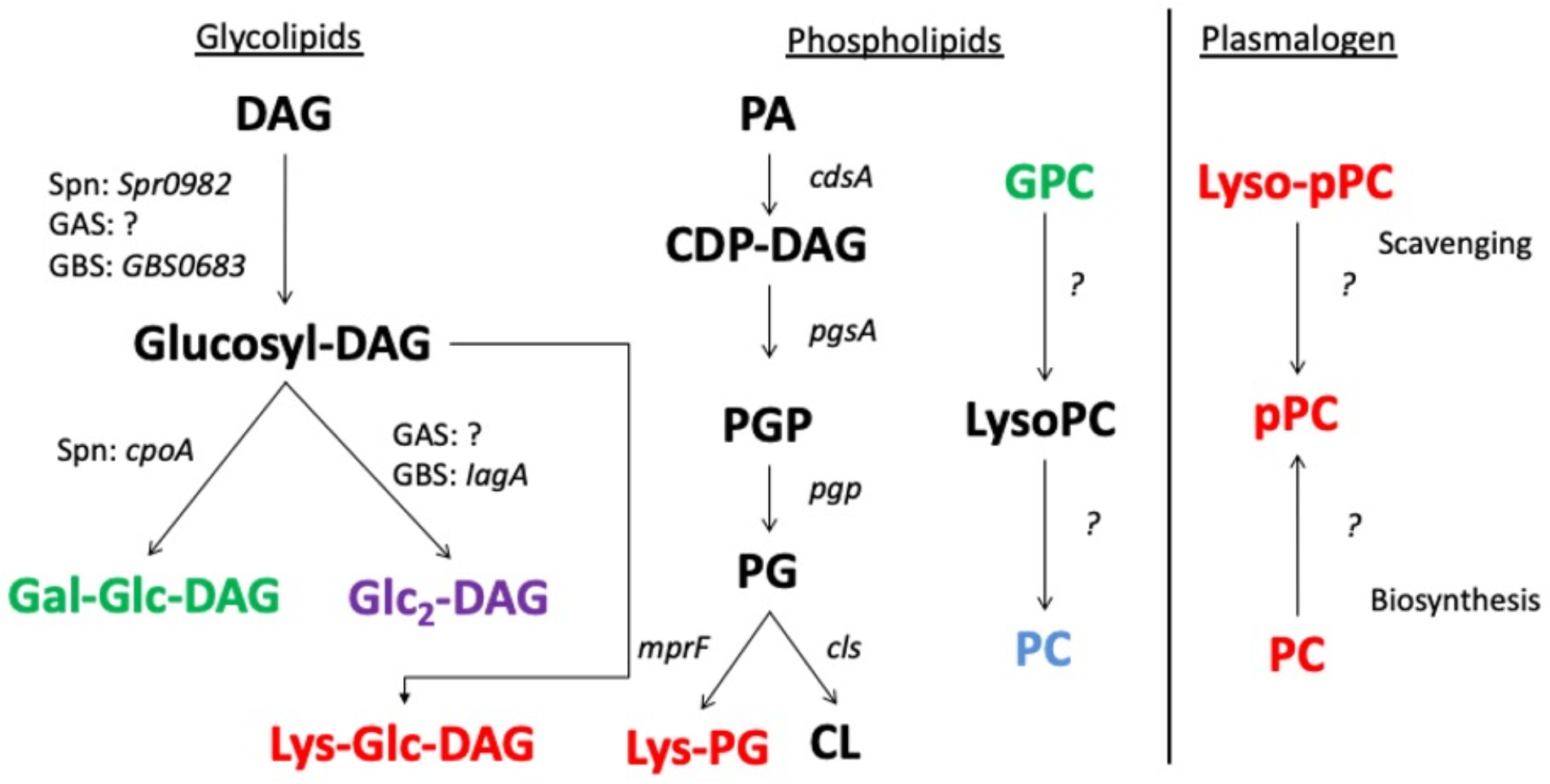
Glycolipid and phospholipid biosynthesis pathways in *S. pneumoniae*, GAS, and GBS. Genes of known or predicted function in each pathway are indicated. Lipids and substrates in black are common to all three species, in green are specific to *S. pneumoniae* (Spn), in red are specific to GBS, in blue are present in *S. pneumoniae* and GAS, and in purple are present in both GAS and GBS. DAG, diacylglycerol; Glc2-DAG, diglucosyl-DAG; Gal-Glc-DAG, galactosyl-glucosyl-DAG; Lys-Glc-DAG, lysyl-glucosyl-DAG (synthesized by *mprF* (23)); PA, phosphatidic acid; CDP-DAG, cytidine diphosphate-DAG; PGP, PG-3-phosphate; PG, phosphatidylglycerol; Lys-PG, lysyl-phosphatidylglycerol; CL, cardiolipin; GPC, glycerophosphocholine; lysoPC, lyso-phosphatidylcholine; PC, phosphatidylcholine; Lyso-pPC, lyso-plasmanyl-PC; pPC, plasmanyl-PC. “?” denotes unidentified genes. It is currently unknown whether GBS scavenge lyso-pPC or pPC from human serum or if it is *de* novo synthesized from PC. This figure combines lipids and genes described in literature (14–19,21–25,32,33) and detected lipids from Table 1.

The lipid profile of *S. pneumoniae* cultured in Todd-Hewitt Broth supplemented with 0.5% w/v yeast extract consists of the major anionic phospholipids PG and CL, the zwitterionic phospholipid PC, and the glycolipids MHDAG and DHDAG (25). Figure 2A (panel 1) displays the positive ESI mass spectrum of PC (appearing at the retention time 19-20.5 min) in the membrane of *S. pneumoniae*. We previously determined that Todd-Hewitt Broth contains GPC, a substrate utilized by *S. pneumoniae* to synthesize PC (25). A similar phospholipid profile is observed for *S. pneumoniae* cultured in defined medium, except that PC is absent, due to the lack of GPC and lysoPC substrates in the medium (25). The total ion chromatogram (TIC) (Supplemental Figure S2) and mass spectrum (MS) of retention time 19-20.5 min of *S. pneumoniae* D39 grown in defined medium lacking human serum are shown in Figure 2A (panel 2), and no PC is present. However, *S. pneumoniae* synthesizes PC when defined medium is supplemented with 5% v/v human serum (Supplemental Figure S2). Figure 2A (panel 3) shows the ESI mass spectrum of the major PC species. These data demonstrate that human serum provides the substrates required for PC biosynthesis via the GPC pathway in *S. pneumoniae*.

**Figure 2.**
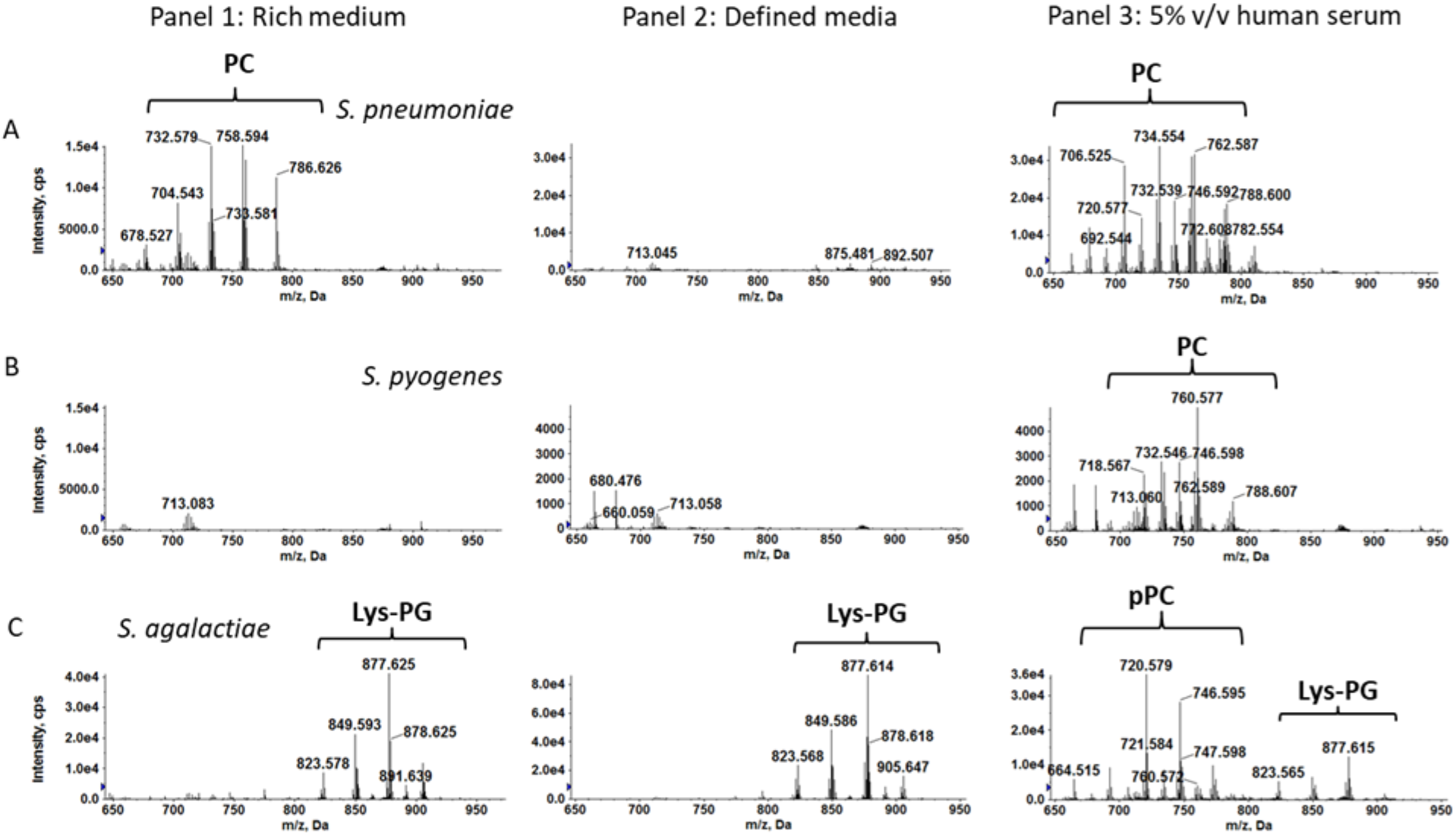
PC and Lys-PG detection when streptococci are cultured in different media. Shown are representative positive ESI mass spectra obtained during the LC retention time of 19 – 20.5 min indicating the presence or absence of PC, pPC, and Lys-PG in the membranes of A) *S. pneumoniae* TIGR4, B) *S. pyogenes* (GAS) MGAS315, and C) *S. agalactiae* (GBS) COH1 when cultured in: Panel 1, rich undefined medium; Panel 2, defined medium; and Panel 3, defined medium supplemented 5% v/v human serum. All cultures were performed in biological triplicate and representative spectra are shown.

The lipid profile of GAS cultured in Todd-Hewitt Broth supplemented with 0.2% w/v yeast extract is similar to that of *S. pneumoniae*, with the exception that PC is not detected (Table 3.1, Figure 2B [panel 1]). The lipid profile remains unchanged in defined medium, and no PC is detected (Figure 2B [panel 2] and Supplemental Figure S2). Interestingly, PC is observed in the membrane of GAS when defined medium is supplemented with 5% v/v human serum, indicating that human serum, but not Todd-Hewitt Broth or defined medium, provide substrates required for PC biosynthesis (Supplemental Figure S2). The ESI/MS of the major PC species identified in the GAS membrane is shown in Supplemental Figure S3. The chemical structure of PC (16:0/18:2) (with the M^+^ ion at *m/z* 758) is shown in Supplemental Figure S3A and the supporting tandem mass spectrometry (MS/MS) fragmentation of *m/z* 758 is shown in Supplemental Figure S3B.

The lipid profile of GBS cultured in Todd-Hewitt Broth consists of MHDAG, DHDAG, PG, and CL, as well as the aminoacylated PG molecule lysyl-PG (Lys-PG), and a recently described novel, cationic glycolipid, Lysyl-Glucosyl-DAG (LGD) (Table 1, Supplemental Figure S2) (23). The lipidomic profile of the GBS is unchanged when cultured in defined medium (Table 1). Like GAS, the GBS do not synthesize PC when cultured in Todd-Hewitt Broth or defined medium (Figure 2C [panel 1 and panel 2]). Also, like GAS, GBS synthesize PC when defined medium is supplemented with 5% v/v human serum (Figure 2C [panel 3], Supplemental Figure S2). However, the ESI mass spectrum (Figure 2C [panel 3]) indicates a modified PC molecule that co-elutes off the column at the same time as Lys-PG. The modified PC molecule was determined to be the plasmalogen, plasmanyl-PC (pPC) (Supplemental Figure S3), in which the *sn*-1 acyl chain linkage is an ether bond, instead of the ester bond that is present in diacyl PC (Supplemental Figure S3A and S3C). The identification of pPC is supported by MS/MS. Specifically, the *m/z* 482 ion in the MS/MS spectrum of *m/z* 720 (Supplemental Figure S3D) is consistent with the pPC structure, as compared to the *m/z* 496 ion in the MS/MS spectrum of *m/z* 758 for the diacyl PC (Supplemental Figure S3B).

Taken together, these data demonstrate that major pathogenic streptococci remodel their membrane lipid composition by synthesizing PC in response to metabolites present in human serum. Furthermore, plasmanyl-PC is uniquely observed in the membrane of the GBS during culture with human serum.

### GAS and GBS scavenge lysoPC to form PC

Next, we sought to investigate if GAS and GBS utilize the GPC biosynthetic pathway previously identified in the Mitis group streptococci (25). GAS and GBS strains were cultured in defined medium supplemented with either 100 μM GPC or 100 μM lysoPC (20:0), a non-natural acyl chain length in these species, and lipidomic analysis was performed. These concentrations are physiologically relevant because lysoPC and GPC are present in varying concentrations throughout the human body at ≤200 μM and ≤500 μM, respectively (30,31). The ESI mass spectra are shown in Figure 3. When GAS is cultured in GPC-supplemented defined medium, very little PC is detected (Figure 3A). In defined medium supplemented with lysoPC (20:0), a substantial amount of PC is detected for GAS (Figure 3B). Similarly, when GBS is cultured in GPC-supplemented defined medium, no PC is detected (Figure 3C), and robust levels of PC are observed when GBS are cultured in defined medium supplemented with lysoPC (20:0) (Figure 3D). Taken together, these data identify lysoPC, not GPC, as the primary substrate scavenged by the GAS and GBS to synthesize PC.

**Figure 3.**
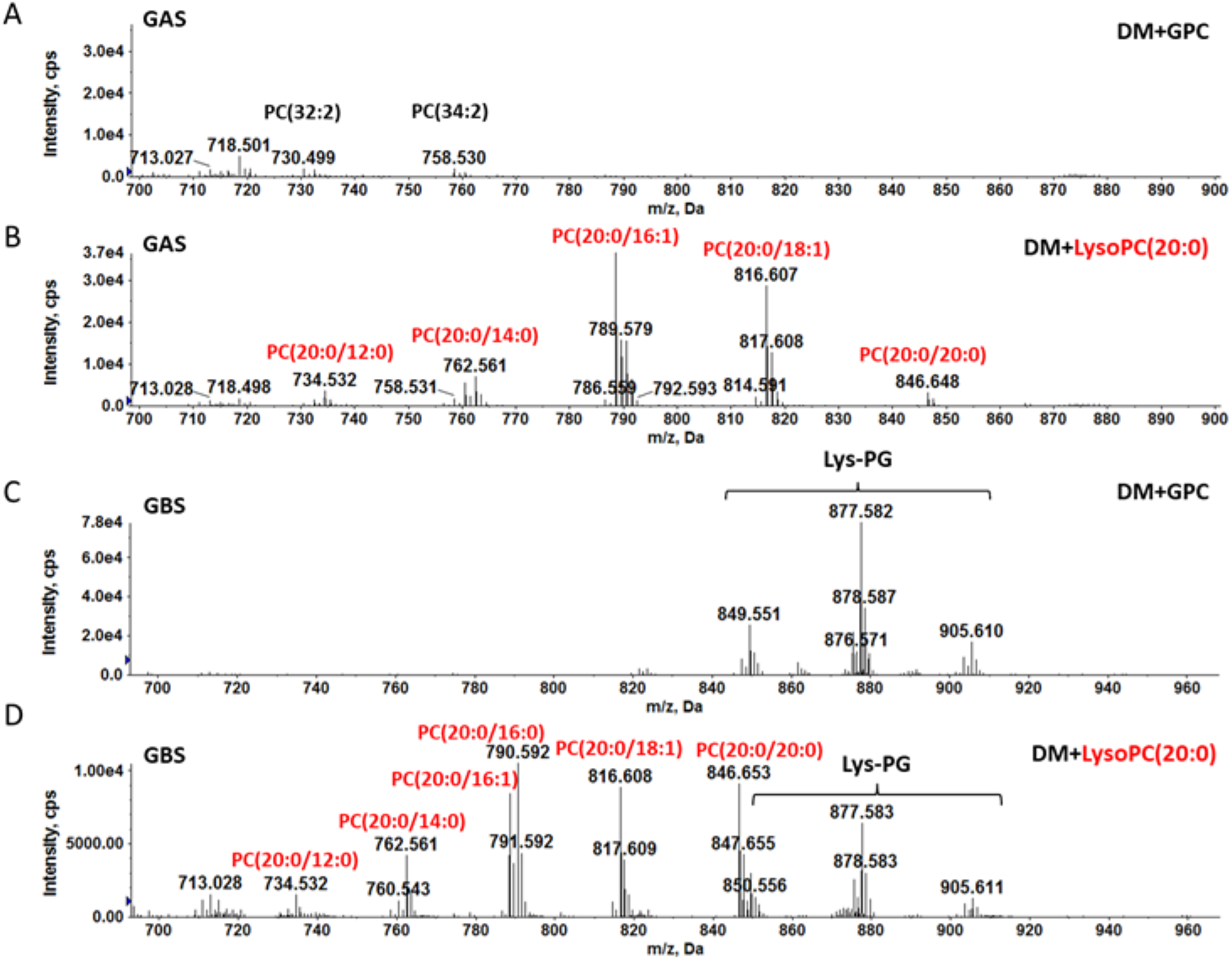
LysoPC is scavenged by GAS and GBS to synthesize PC. A) GAS strain NZ131 cultured in defined medium supplemented with 100 μM GPC synthesizes a very low level of PC. B) GAS strain NZ131 grown in defined medium supplemented with 100 μM lysoPC (20:0) synthesizes PC. C) No PC is detected for GBS strain COH1 cultured in defined medium supplemented with 100 μM GPC. D) GBS strain COH1 cultured in defined medium supplemented with 100 μM lysoPC (20:0) synthesizes PC. Cultures were performed in biological triplicate and representative data are shown.

## Discussion

The cellular membrane is a critical and dynamic surface of bacterial cells that plays a role in many cellular processes yet is largely understudied in streptococci. In this study we characterized the lipidome of the major streptococcal pathogens *S. pneumoniae*, GAS, and GBS when cultured in a standard rich laboratory medium (Todd-Hewitt Broth), and a defined medium supplemented with or without 5% v/v human serum. We show that all three streptococcal species remodel their cellular membrane in response to human metabolites, specifically, by synthesis of PC. To our knowledge, this is the first identification of PC and the plasmalogen pPC in the GAS and GBS, respectively. PC has been linked to virulence in certain bacteria (32,33). Streptococci may incorporate PC into their membranes as a form of eukaryotic membrane mimicry to evade immune defense, although this must be investigated further.

Human serum is a rich source of nutrients for streptococci. A major metabolite of human serum is lysoPC (30,34). *S. pneumoniae* utilizes a full GPC pathway (25), scavenging metabolites from human serum to synthesize PC. However, the GAS and GBS predominantly synthesize PC using an abbreviated GPC pathway, by primarily scavenging lysoPC from the exogenous environment. This is likely due to a missing or poorly expressed GPC transporter and/or the inability to acylate GPC to form lysoPC. The ability of *S. pneumoniae* but not GAS or GBS to scavenge GPC could be capitalized upon in comparative genomic and laboratory experiments to help identify the GPC transporter in *S. pneumoniae*, which is currently unknown.

To our knowledge, the presence of the plasmalogen pPC in the GBS membrane has not been described before. To date, plasmalogens have only been observed in strict anaerobes such as *Clostridium* and *Bifidobacterium* (35–37). Biosynthesis of plasmalogens in bacteria is poorly understood, with the first bacterial desaturase involved in plasmenyl *sn*-1 vinyl ether bond formation and the first plasmalogen synthase, PlsAR, in *Clostridium perfringens* only recently identified (38,39). Plasmalogen lipids are thought to promote oxidative stress survival in animal cells (40,41) and may play a role in GBS pathogenesis. pPC has been detected in human blood at ~50 μM in healthy adults (30,34). There are two possible pathways for the presence of pPC in the membrane of GBS during culturing with human serum. Either, 1) GBS scavenge the intermediate lyso-pPC, adding the second acyl chain, and/or full pPC directly from human serum or 2) *de novo* pPC synthesis is occurring. Our experimental design does not allow us to discriminate which pathway is utilized. Given that GBS lack an identifiable homolog of the *C. perfringens* PlsAR synthase, coupled with the lack of plasmalogen form of other common lipids, the pPC we identify in the membrane of GBS is likely derived from the plasmalogen lipids present in human serum. The origin and roles of pPC in GBS is a subject of further investigation.

Overall, this work demonstrates that culture medium can significantly alter the membrane lipid composition of major human streptococcal pathogens. This supports the significance of culturing pathogens in media that more closely represent the host environments in which the pathogens are found. Further investigation into the biosynthesis and mechanistic roles of PC in the membrane will likely provide novel insights into the pathogen-host interactions and pathogenesis of these streptococcal species.

## Acknowledgements

We gratefully acknowledge Michael Federle at the University of Illinois at Chicago for providing *S. pneumoniae* D39.

## Supplemental Text S1. Materials and Methods

### Growth curves

GAS and GBS were cultured overnight in defined medium and diluted to a starting OD_600nm_ of 0.05 in 15 mL pre-warmed defined medium supplemented with or without 5% v/v human serum. The OD_600nm_ was monitored manually every hour using a Thermo Scientific Genesys 30 spectrophotometer. For *S. pneumoniae*, early exponential phase defined medium overnight cultures, as described above, were diluted 1:50 into pre-warmed defined medium supplemented with or without 5% v/v human serum. *S. pneumoniae* cultures were incubated for 4 h before the OD_600nm_ was monitored every hour. Growth curves were performed in biological triplicate. Repeated measures two-way ANOVA with Bonferroni’s multiple comparisons test was performed in GraphPad Prism version 8 for Windows, GraphPad software, San Diego, California, USA, www.graphpad.com.

### Acidic Bligh-Dyer extractions

Centrifugation was performed using a Sorvall RC6+ centrifuge. Cultures were pelleted at 4,280 x *g* for 5 min at room temperature. The supernatants were removed and stored at −80°C until acidic Bligh-Dyer lipid extractions were performed as described (1,2). Briefly, cell pellets were resuspended in 1X PBS (Sigma-Aldrich) and transferred to Coring Pyrex glass tubes with PTFR-lined caps (VWR), followed by 1:2 vol:vol chloroform:methanol addition. Single phase extractions were vortexed periodically and incubated at room temperature for 15 minutes before 500 x *g* centrifugation for 10 min. A two-phase Bligh-Dyer system was achieved by addition of 100 μl 37% HCL, 1 mL CHCl_3_, and 900 μl of 1X PBS, which was then vortexed and centrifuged for 5 min at 500 x *g.* The lower phase was removed to a new tube and dried under nitrogen before being stored at −80°C prior to lipidomic analysis.

### Normal-Phase Liquid Chromatography-Electrospray Ionization/Mass Spectrometry (LC-ESI/MS)

Normal-phase LC-ESI/MS was performed on an Agilent 1200 quaternary LC system equipped with an Ascentis silica high-performance liquid chromatography (HPLC) column (5 μm; 25 cm by 2.1 mm; Sigma-Aldrich) as described previously (1–3). Briefly, mobile phase A consisted of chloroform-methanol-aqueous ammonium hydroxide (800:195:5, vol/vol), mobile phase B consisted of chloroform-methanol-water-aqueous ammonium hydroxide (600: 340:50:5, vol/vol), and mobile phase C consisted of chloroform-methanol-water-aqueous ammonium hydroxide (450:450:95:5, vol/vol/vol/vol). The elution program consisted of the following: 100% mobile phase A was held isocratically for 2 min, then linearly increased to 100% mobile phase B over 14 min, and held at 100% mobile phase B for 11 min. The LC gradient was then changed to 100% mobile phase C over 3 min, held at 100% mobile phase C for 3 min, and, finally, returned to 100% mobile phase A over 0.5 min and held at 100% mobile phase A for 5 min. The LC eluent (with a total flow rate of 300 ml/min) was introduced into the ESI source of a high-resolution TripleTOF5600 mass spectrometer (Sciex, Framingham, MA). Instrumental settings for positive-ion ESI and MS/MS analysis of lipid species were as follows: IS = 5,000 V, CUR = 20 psi, GSI = 20 psi, DP = +55 V, and FP = +150V. The MS/MS analysis used nitrogen as the collision gas. Data analysis was performed using Analyst TF1.5 software (Sciex, Framingham, MA).

**Supplemental Figure S1.**
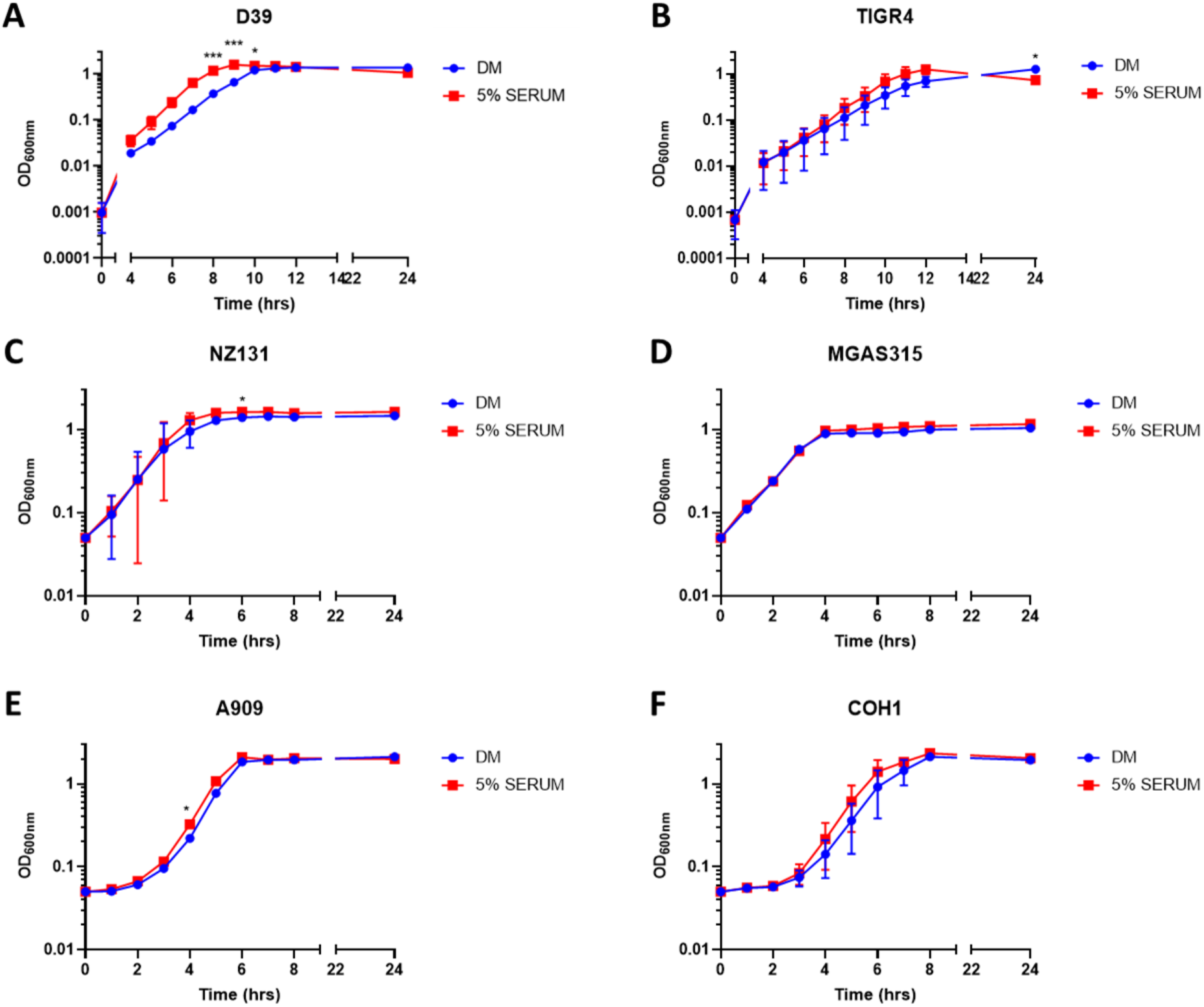
Growth characteristics of streptococci in the presence of 5% human serum. Growth curves in defined medium (DM) are shown in blue and defined medium supplemented 5% v/v human serum are shown in red. A) *S. pneumoniae* D39, B) *S. pneumoniae* TIGR4, C) *S. pyogenes* NZ131, D) *S. pyogenes* MGAS315, E) *S. agalactiae* A909, and F) *S. agalactiae* COH1. Manual OD_600nm_ readings were performed every hour. *S. pneumoniae* cultures were grown for 4 h before manual readings were performed. Growth curves were performed in biological triplicate. Mean and SD are indicated. Statistical analysis: A-F) repeated measures two-way ANOVA with Bonferroni’s multiple comparisons test. * denotes p-value <0.05. *** denotes p-value <0.001.

**Supplemental Figure S2.**
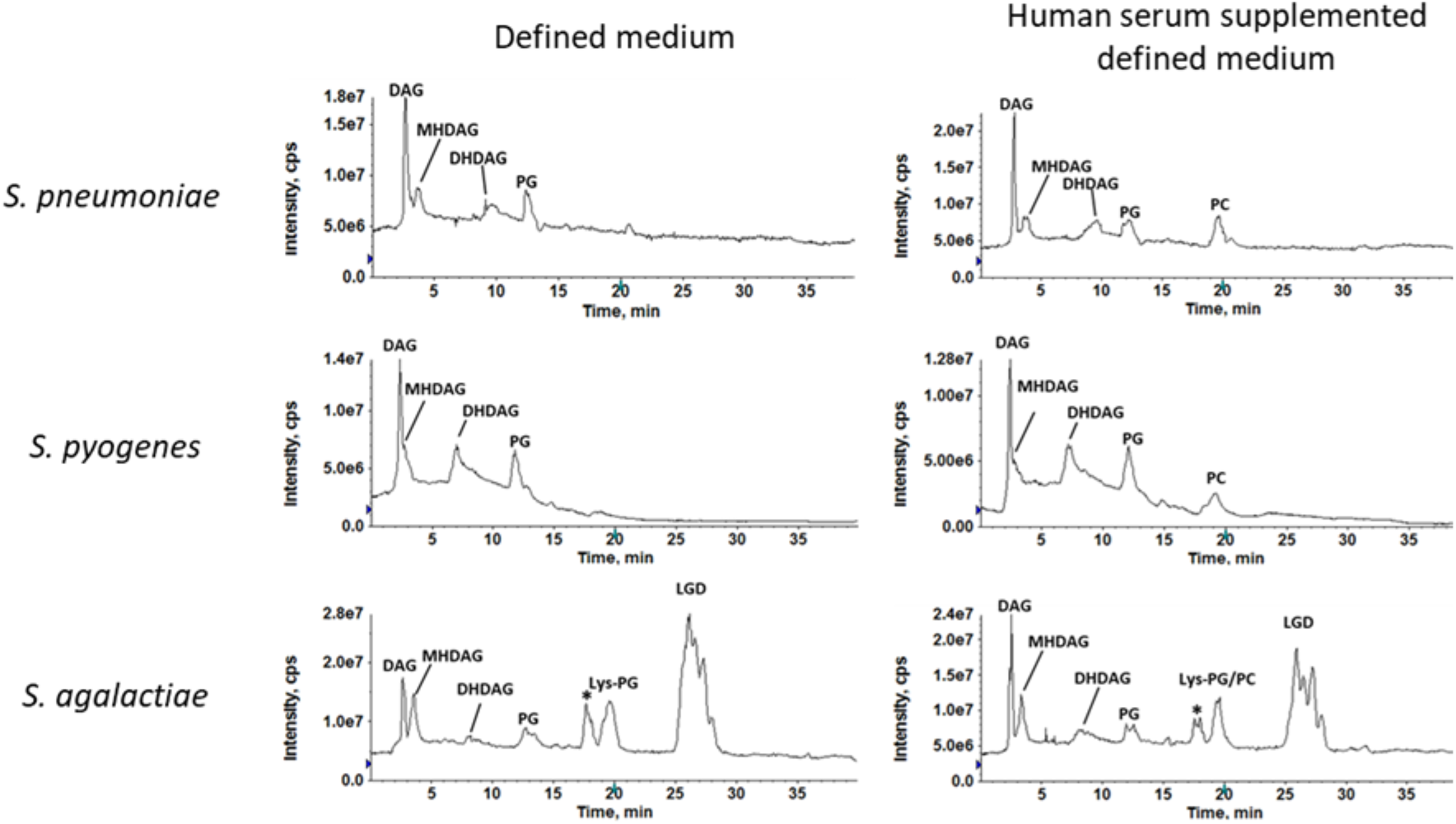
Positive ion mode total ion chromatograms (TIC) of streptococci grown in defined medium supplemented with or without 5% v/v human serum. Representative positive ion mode TIC for *S. pneumoniae* (top panels), *S. pyogenes* (middle panels), and *S. agalactiae* (bottom panels), indicate relative abundance of lipids present in the cell extract when streptococci are cultured in defined medium or defined medium supplemented with 5% v/v human serum. Cultures were performed in biological triplicate. Abbreviations: DAG, diacylglycerol; MHDAG, monohexosyldiacylglycerol; DHDAG, dihexosyldiacylglycerol; PG, phosphatidylglycerol; Lys-PG, lysyl-phosphatidylglycerol; LGD, lysyl-glucosyl-DAG; PC, phosphatidylcholine; ^‘^*^’^ Denotes methylcarbamate, an extraction artifact of LGD.

**Supplemental Figure S3.**
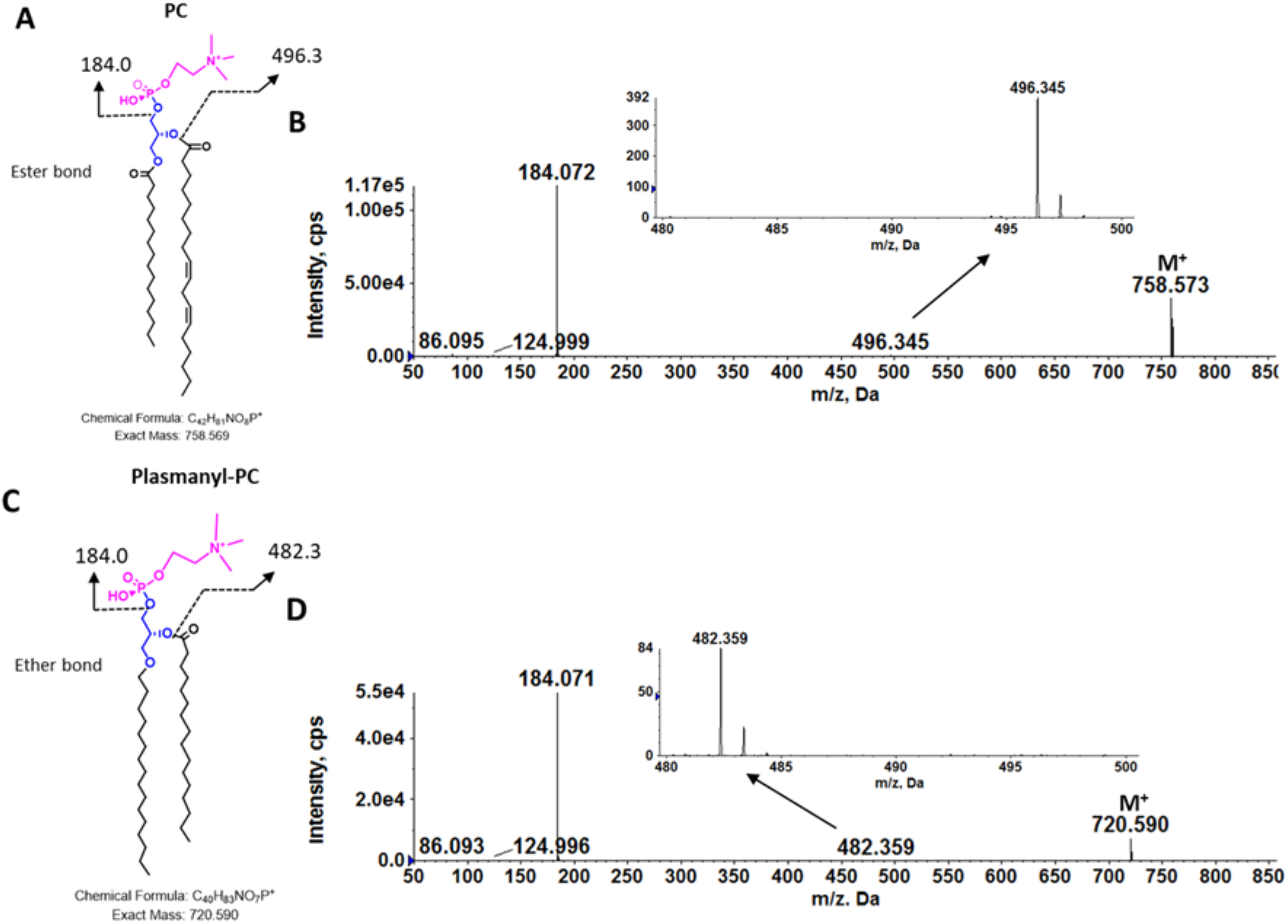
Confirmation of major PC and plasmanyl-PC species by MS/MS fragmentation. A) Chemical structure and fragmentation scheme of PC(16:0/18:2), B) MS/MS of *m/z* 758 for PC(16:0/18:2), C) Chemical structure and fragmentation scheme of pPC(16:0/16:0), and D) MS/MS of *m/z* 720 for pPC(16:0/16:0). MS/MS fragmentation, specifically *m/z* 482 fragment, is consistent with the structure of pPC present in the GBS.

**Table S1.**
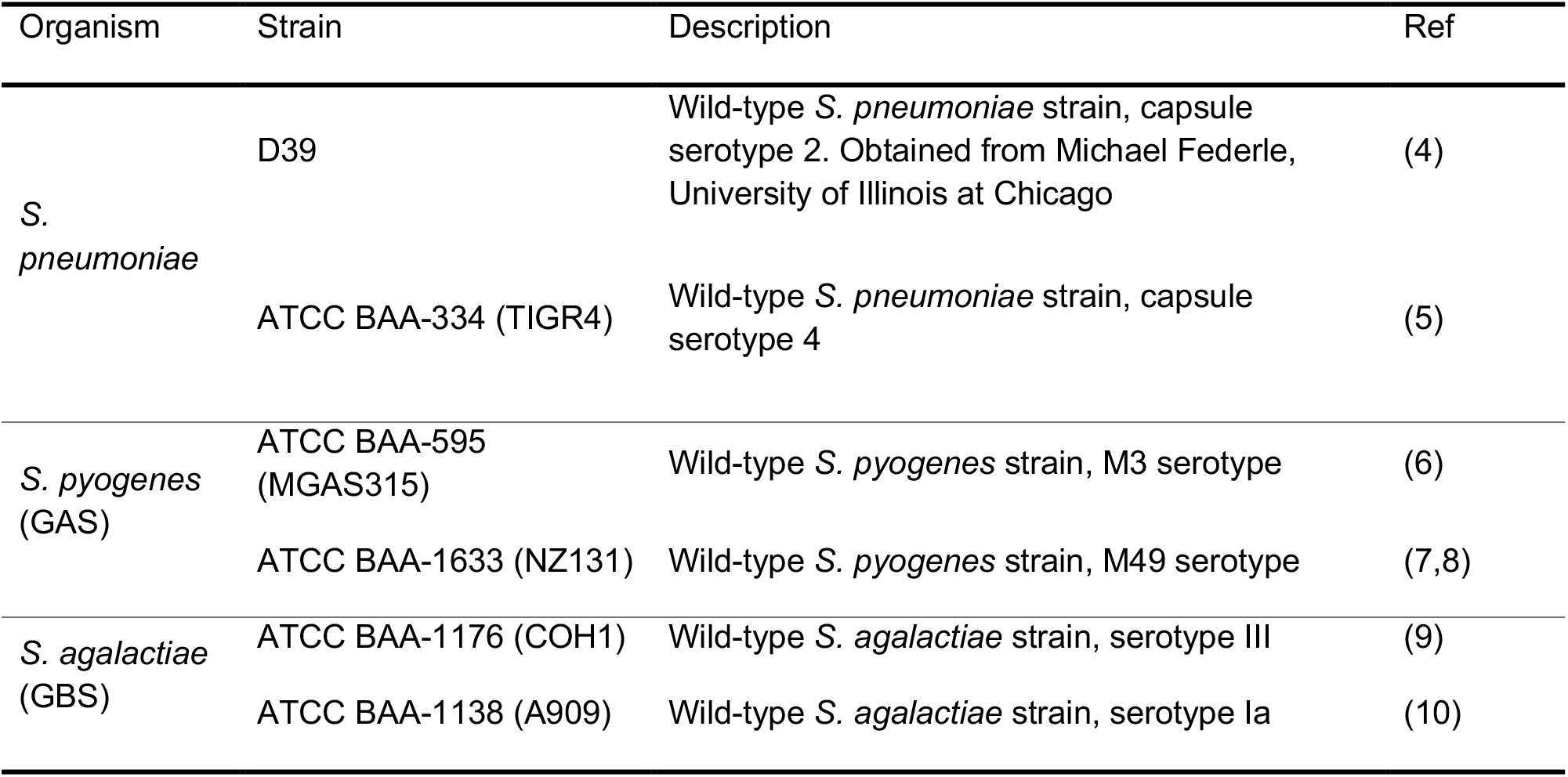
Strains used in this study.

